# Understanding the role of c-di-AMP signaling in determining antibiotic tolerance in *Mycobacterium smegmatis*: generation of resistant mutants and regrowth of persisters

**DOI:** 10.1101/2022.05.06.490884

**Authors:** Aditya Kumar Pal, Anirban Ghosh

## Abstract

In this study, we probe the role of secondary messenger c-di-AMP in drug tolerance, which includes both persister and resistant mutant characterization of *Mycobacterium smegmatis*. Specifically, with the use of c-di-AMP null and overproducing mutants, we showed how c-di-AMP plays a significant role in resistance mutagenesis against antibiotics with different mechanisms of action. We elucidated the specific molecular mechanism linking the elevated intracellular c-di-AMP level and high mutant generation and highlighted the significance of non-homology-based DNA repair. Further investigation enabled us to identify the unique mutational landscape of target and non-target mutation categories linked to intracellular c-di-AMP levels. Overall fitness cost of unique target mutations was estimated in different strain backgrounds, and then we showed the critical role of c-di-AMP in driving epistatic interactions between resistance genes, resulting in the evolution of multi-drug tolerance. Finally, we identified the role of c-di-AMP in persister cells regrowth and mutant enrichment upon cessation of antibiotic treatment.

## Introduction

Bacterial second messengers are a group of nucleotide-derived small molecules that modulate important cellular pathways in different bacteria. A few examples of those messenger molecules are (p)ppGpp, c-di-GMP, c-di-AMP, cGAMP etc. ^1–6^. So far, especially in Gram-positive organisms, c-di-AMP has been shown to contribute to different biological processes such as DNA repair, K^+^ transport, osmotic balance maintenance, cell wall metabolism, virulence etc. ^7–11^. In some cases, it was also studied to induce the type-I interferon response in mammalian cells ^12,13^. Our group was specifically interested to study if c-di-AMP could play an important role in terms of promoting antibiotic tolerance in mycobacteria. Though there are few studies of c-di-AMP in *M. smegmatis* and *M. tuberculosis* ^14–19^ which mainly highlighted key pathways regulated by c-di-AMP, there was no study that exclusively described the physiological role of this important small molecule to promote resistant mutant generation in any bacteria. Since one of our recent studies identified a potential impact of varying intracellular c-di-AMP concentration regulating distinctive drug susceptibility phenotypes in *M. smegmatis* ^20^, we wanted to continue our research in the same direction to understand such differential drug tolerance mechanisms and increased probability of resistant mutant generation. In *Mycobacterium smegmatis*, c-di-AMP is constitutively synthesized from the condensation of two ATP molecules by enzyme DisA and hydrolyzed by enzyme Pde into pApA^14^; often this steady-state homeostasis gets unbalanced as a part of stress adaptation. As a matter of course, further understanding was demanded if and how the varying c-di-AMP concentration could play a role in the antibiotic resistance phenotype of the cells and our data indeed highlighted the strong correlation between intracellular c-di-AMP concentration and the evolution of resistant mutants. We elucidated the mechanistic insights and our model suggests the role of c-di-AMP concentration in determining specific categories of mutations in different growth phases. Next, we revealed the unexpected role of c-di-AMP in controlling epistatic interaction between two resistance genes of mycobacteria and thus stimulating multi-drug resistance phenotype. Finally, we show how c-di-AMP contributes to the resuscitation of persisters by transcriptional regulation of resuscitation-promoting factors through a putative riboswitch element. All in all, this study discusses the physiological relevance of c-di-AMP in modulating the antibiotic resistance profile of *M. smegmatis* and possible implications in the evolution of persisters and resistant mutants.

## Materials and Methods

### Bacterial strains, media and growth conditions

*M. smegmatis* mc^2^155 (WT) and its knockout variants *ΔdisA & Δpde* and their respective complemented strains (*ΔdisA+pDisA*) & (*Δpde+pPde*) were grown in Middlebrook 7H9 broth (MB7H9; HiMedia) with 2% (wt/vol) glucose as a carbon source and 0.05% (vol/vol) tween-80, Agar (1.5%, w/v) (HiMedia) at 37°C. The antibiotics kanamycin and hygromycin were used at a concentration of 25 μg/ml and 50 μg/ml respectively ^20^. ciprofloxacin, rifampicin and Ofloxacin powder were obtained from Sisco Research Laboratories. Antibiotics were used at variable concentrations.

### Estimation of resistant mutation frequencies and rates

As previously described ^21^, single colonies from *M. smegmatis* strains were grown in Middlebrook 7H9 medium to an optical density (OD at 600nm, OD600) of 1.2-1.5. Next, 1% of the inoculum had given to a fresh 10 ml. of media and kept at 37°C shaking. 1ml. of cultures of each strain was taken from exponential (OD_600_ 0.7-0.8) and stationary (OD_600_ 2.5-3) phase cultures and concentrated by 5 times. 100 μl of the culture was plated in triplicate on MB agar plates containing antibiotics to get the mutant cell count and the remaining cultures were diluted till 10^-7^ dilution and 100ul of selective dilutions were plated in triplicate onto an antibiotic-free MB agar for CFU determination (drug-free control). Mutation frequencies were calculated by dividing the number of colonies on a drug-containing plate by the number of colonies on the drug-free plate ^22^. In addition, 50-100 colonies were chosen randomly from the drug-containing plates for further characterization.

Spontaneous mutation rates of *M. smegmatis* strains to ciprofloxacin and rifampicin were determined by the Luria-Delbrück fluctuation analysis using the method described by ^23^ with minor modification. Single colonies of *M. smegmatis* wild-type, *ΔdisA* and *Δpde* strains were grown in Middlebrook 7H9 medium to OD_600_ of 1.2-1.5 and for each strain, 1000-1500 cells/ml of inoculum had given into 10 tubes and kept at 37°C shaking for 6 days. For selecting the spontaneous mutants, the whole culture was concentrated by 3 times and plated on drug-containing plate and 50 μl was kept aside for the CFU estimation. Mutation rates were calculated by the Luria and Delbruck formula ^24^ **(Fig. S1d)**:

### Ultra-violet stress induced mutation frequency

*M. smegmatis* strains were grown until the mid-log phase and then normalized to OD_600_ 0.7. 1 ml. of this culture was exposed to UV irradiation (0.125 mJ/ cm^2^) ^25^. After that, an inoculation was given from the UV treated and untreated cultures into fresh Middlebrook 7H9 broth to make 0.03 OD and kept at 37°C shaking for 5 days. On the fifth day, the 100 μl of the cultures were spread into rifampicin plates (10X MIC concentration) to estimate the resistant mutant population and also 50 μl of culture was taken for the CFU estimation in an antibiotic-free plate. Mutation frequency was determined using the formula discussed in the mutation frequency method section.

### Plasmid based NHEJ *in-vivo* assay

The non-homologous end-joining (NHEJ) reporter plasmid *pMV261-lacZ* (hygromycin marker) was constructed by cloning a functional copy of the *lacZ* gene from the pSD5B vector using the HindIII restriction site in specific primers (Table S7). For *in vivo* NHEJ reporter assay, we adapted the protocol from a previously published paper ^26^. In our case, the pMV261-*lacZ* reporter vector was digested with blunt cutter restriction enzyme SspI. The linear plasmid was purified by agarose gel electrophoresis and the DNA concentration was quantified by UV absorbance. The same concentration of linear and circular DNAs (uncut pMV261-*lacZ* plasmid) was transformed into different *M. smegmatis* strains by electroporation and plated on Middlebrook agar containing hygromycin (50μg/ml) and X-gal (40μg/ml). Transformations were done in triplicates for each strain and the plates were kept at 37°C for 72 hours. Colonies were counted after 72 hours. The blue colony colour indicated an intact *lacZ* coding sequence; whereas the white colonies indicated that the *lacZ* gene was inactivated during the repair process. NHEJ efficiency was calculated by the ratio of CFU per nanogram of transformed linear DNA and circular DNA. NHEJ fidelity was calculated as the percentage of blue colonies out of the total number of colonies ^27^. A few white mutant colonies of *Δpde* pMV261-*lacZ* were isolated from the plate and streaked onto a new X-gal containing plate to reconfirm the loss of *lacZ* function. Colony PCR was performed by taking one single colony dissolved in 50μl of autoclaved water. PCR products were isolated by using the Qiaquick PCR cleanup kit (Qiagen) and then sequenced directly using the lacZ_MIDREV2 primer (Table S7) to detect mutations.

### Minimum Inhibition Concentration (REMA) assay

MIC values were estimated using REMA (Resazurin microtiter assay) adapted from an earlier protocol ^28^. In brief, the transmittance of the culture was adjusted to a McFarland turbidity standard of 1 and then diluted to 1:10. Next, 196 μl portions of the diluted culture were inoculated into 96 well microtiter plates containing a 2-fold serial dilution of the antibiotics (4 μl). Plates were sealed and incubated at 37°C for 36 hours. Next, 30 μl of 0.01% resazurin dye was added to each well, and the plates were further incubated for 4-6 hours. The color of the resazurin changed from blue to pink due to bacterial growth. The MIC was determined as the minimum antibiotic concentration at which the resazurin dye did not change the color.

### Mutation mapping of cip^R^ and rif^R^ mutants

Antibiotic-resistant mutant colonies were isolated from different concentrations of drug-containing plates, inoculated into the same antibiotic containing Middlebrook 7H9 complete media and were grown in shaking at 37°C, till the mid-log phase (MLP). The genomic DNA of the mutants was isolated using the phenol-chloroform extraction method ^29^ after the overnight lysis procedure. The quinolone resistance determining region (QRDR) of *gyrA* and the rifampicin resistance determining region (RRDR) of the *rpoB* gene were amplified from the genomic DNA of respective resistant mutants using the specific primer pairs (Table S7). The sequencing reactions were performed by Medauxin.

### EtBr efflux Assay

Efflux assay was adapted from ^30^ a previous publication. Briefly, *M. smegmatis* cultures were grown at 37°C in MB7H9 medium to an optical density at 600nm of 0.9-1. Cells were then washed twice with phosphate buffer saline with 0.05% tween80, resuspended in 1/3^rd^ volume of the same buffer kept at 37°C shaking for one hour for starvation. After that, CCCP (carbonyl cyanide m-chlorophenylhydrazone) (100 μM) was added (acts as an efflux pump inhibitor ^31^ and the cells were further incubated for 30 minutes in the same condition. Next, Ethidium bromide (0.5 μg/ml f.c.) was added and cells were incubated for another 30 minutes. After that, cells were washed with the 1XPBST in the same volume to remove the CCCP and extracellular EtBr. After that, cells were taken into 96 well black plates and 2% glucose was added to the wells to facilitate efflux activity. To quantify the efflux of EtBr fluorescence was checked for the next one hour (with a 1-minute interval between two readings) in the Varioskan Flash multimode reader (Thermo Fisher Scientific) at 530 nm and 590 nm wavelengths for excitation and emission respectively.

### Fitness cost estimation of mutants and analysis of epistatic interaction

Ciprofloxacin resistant (cip^R^) mutants were grown in MB7H9 media till the mid-exponential phase, washed and inoculated into M9 minimal media containing 1mM MgCl2, 0.3mM Cacl2 with 0.2% glucose ^32^ by adjusting the final OD_600_ 0.03. 5 ml. of such cultures were taken in a sterile glass tube and kept at 37°C shaking for 120 hours and OD_600_ was measured in equal intervals. To check if the fitness cost of certain cip^R^ mutants was reversed by a secondary mutation and emergence of rifampicin resistance, cells were plated on rifampicin (10X MIC) plates after 120 hours of growth in M9 minimal media, and the mutation rate was calculated.

### Regrowth and Resuscitation assay with mutant enrichment determination

*M. smegmatis* Cultures were grown till the late exponential/early stationary phase (OD_600_ of 1.5-2.5), diluted in 1:100 to start a secondary culture, and grown till the mid-exponential phase (OD_600_ of 0.6-1) and further diluted to adjust to OD_600_ of 0.2 corresponding to ~2 X10^7^ CFU/ml (CFU= colony-forming unit) in fresh MB7H9 medium. 5 ml of such culture was taken for ciprofloxacin (3X MIC) treatment at 37°C shakers. CFU estimations (plating) were done every 12 hours for up to 72 hours. In parallel to the CFU estimation, spotting/spreading was done in ciprofloxacin (1.25 μg/ml.) plates to enumerate resistant mutant population enrichment over time. In the case of resuscitation assay (10X ciprofloxacin), after 24 hours of treatment, cultures were washed to remove antibiotics and resuspended in fresh MB7H9 media without antibiotics followed by incubation again incubated at 37°C shaker and CFU estimations (plating) were done every 24 hours of up to 96 hours. Plates were further incubated at 37°C for 3-4 days for colony growth and subsequently counted.

### GFP expression measurement of the promoter fusion construct

The *rpfA* gene reporter constructs were made in promoter less GFP vector pMN406-Δ*imyc* ^33^ (a generous gift from Prof. Ajitkumar, IISc, India). To generate transcriptional fusions, a 434 bp fragment from the upstream region of the gene *MSMEG_5700* was PCR amplified from *M. smegmatis* mc^2^155 gDNA using a compatible set of primers, as mentioned in (Table S7). The PCR fragment was then subsequently cloned into the reporter vector upstream of GFP using restriction enzymes and transformed into Wild-type, *Δdisa* and *Δpde* strains and the transformants were selected against hygromycin. To measure the GFP induction during the resuscitation phase in different strains, 200 μl of the cells were taken in a black, flat bottom 96 well plate (Thermo Nunc) and GFP intensity was recorded at excitation of 488nm and emission of 510 nm ^34^ in Varioskan Flash multimode reader (Thermo Fisher Scientific). GFP expression (RFU) was calculated by dividing GFP fluorescence intensity by cell density (OD 600nm).

## Results

### High c-di-AMP concentration promotes the generation of spontaneous resistant mutants

We designed our assay to evaluate the resistant mutant frequency of *M. smegmatis* WT, *ΔdisA* and *Δpde* strains in different growth phases, against the well-known fluoroquinolone antibiotic ciprofloxacin. *M. smegmatis* cells were grown till the early stationary phase and spread on high concentration (10X MIC) of ciprofloxacin plates and the number of resistant mutants was counted **(Fig.S1a)**. It was found that the *in vitro*, spontaneous mutants of *M. smegmatis Δpde* resistant to ciprofloxacin arose at frequencies of 7.9 × 10^-7^ whereas for *M. smegmatis* WT strain’s resistance frequency remained low ~4.5 × 10^-8^. This ~18-fold increase in resistance frequency was directly correlated to high c-di-AMP concentration as the complementation strain*M. smegmatisΔpde+* pMV361-*pde* had a similar resistance frequency to WT ~5.2 × 10^-8^ **(Fig.1A)**. For *M. smegmatis ΔdisA* strain there was no significant increase in resistance frequency remaining at 8.3 × 10^-8^ and similar values were observed with complementation strain *M. smegmatis ΔdisA+* pMV361-*disA* **(Fig.1A)**. Next, we wanted to check if this c-di-AMP driven resistance phenotype is specific for well-grown stationary phase cultures only or not, and thus repeated the assay with exponentially grown cultures of the same strains **(Fig.S1a)** and we found that the *M. smegmatis Δpde* strain had ~5 fold high resistance frequency compared to WT, whereas *M. smegmatis ΔdisA* strain had similar resistance profile like WT **(Fig.S1b)**. As expected, both the complementation strain behaved like WT further corroborating the relevance of high c-di-AMP concentration with higher mutation frequency across all growth phases. In general, all the strains grown till the stationary phase showed higher mutation frequency compared to the respective exponentially grown cultures under an identical assay setup. To find out, if the hypermutant phenotype of *M. smegmatis Δpde* strain was not specific to DNA damaging antibiotic ciprofloxacin, we checked the resistance frequency of the strains against RNA polymerase inhibitor rifampicin and we noted a significant ~44-fold increase in resistance frequency for *Δpde* strain. But, to our surprise, *ΔdisA* strain also showed a ~29-fold increase in resistance frequency compared to WT, unlike ciprofloxacin. As expected, both the complementation strains had similar RF values to WT **(Fig.1B, S1c)**.

**Figure 1.**
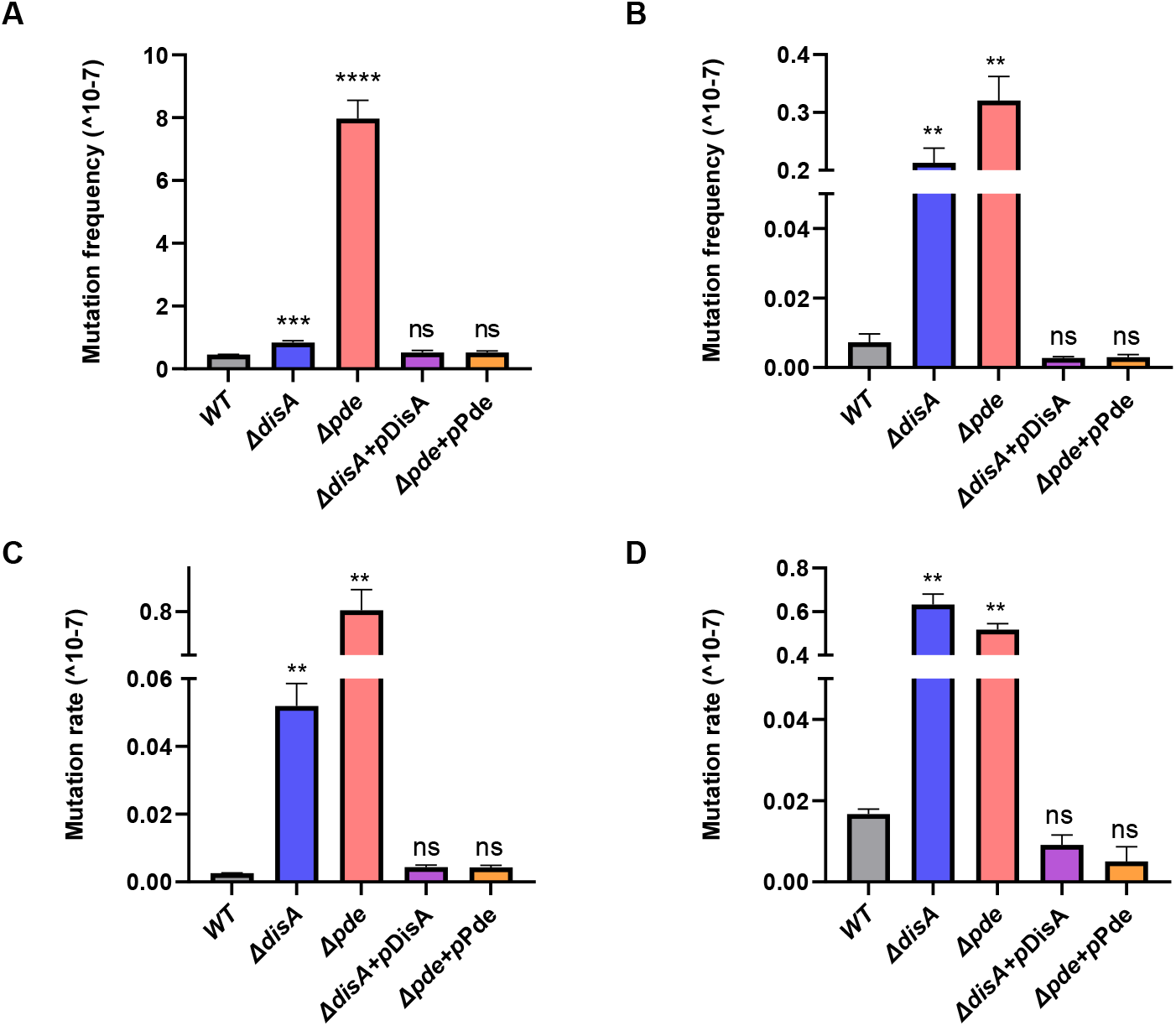
Modulating intracellular c-di-AMP concentration affects the number of spontaneous resistant mutants against ciprofloxacin and rifampicin. **(A)** ciprofloxacin mutation frequency and **(B)** rifampicin mutation frequency were calculated for *M. smegmatis* WT, *M. smegmatis ΔdisA, M. smegmatis Δpde* and their respective complementation strains [N=3]. Similarly, **(C)** ciprofloxacin mutation rate and **(D)** rifampicin mutation rate were calculated for *M. smegmatis* WT, *M. smegmatis ΔdisA*, *M. smegmatis Δpde* along with respective complementation strains [N=2]. All the graphs are plotted using GraphPad Prism8, unpaired t-test was used to calculate statistical significance: *** = P < 0.001; ** = P < 0.01; * = P < 0.05; ns= non-significant.

Spontaneous mutation rates were estimated **(Fig.S1d)** against ciprofloxacin and rifampicin for all 5 strains and a similar observation in terms of increase in mutation rate was observed in *Δpde* strain, which further confirmed the direct role of c-di-AMP in promoting antibiotic resistance **(Fig.1C)**. As a similar observation with mutation frequency assay with rifampicin, here also *M. smegmatis ΔdisA* showed a significant increase in mutation rate **(Fig.1D)**. The specific fold change in spontaneous mutation rates of the strains was illustrated in **Table 1**. At this point, we speculated whether the selective increase in the probability of spontaneous rifampicin mutation of *ΔdisA* strain was due to the lack of the DisA scanning enzyme or c-di-AMP messenger. Hence, we repeated the assay *ΔdisA* strain with complementation of a *disA* mutant (D84A) incapable of synthesizing c-di-AMP ^35^ but unaffected in DNA binding. Our data implied a significant drop in mutation rate in *ΔdisA* strain when complemented with pMV361-*disA*(D84A), similar to pMV361-*disA*(WT) complementation **(Fig.S1e)**. This observation led us to believe that, due to the lack of DisA enzyme’s DNA scanning properties (and not due to the absence of c-di-AMP) in *ΔdisA* strain significant increase in mutation rate was observed.

**Table 1.** Specific fold changes in spontaneous mutation rates are mentioned for different strains against ciprofloxacin and rifampicin.

| Strains | Ciprofloxacin |  | Rifampicin |  |
| --- | --- | --- | --- | --- |
| | Mutation rate<br>($\times 10^{-7}$) | Fold<br>increase | Mutation rate<br>($\times 10^{-7}$) | Fold<br>increase |
| *WT* | 0.002 ± 0.000 | **1** | 0.016 ± 0.001 | **1** |
| *ΔdisA* | 0.051 ± 0.005 | **25.5** | 0.633 ± 0.033 | **39.56** |
| *Δpde* | 0.802 ± 0.033 | **401** | 0.517 ± 0.019 | **32.3** |
| *ΔdisA+pDisA* | 0.004 ± 0.000 | **2** | 0.009 ± 0.001 | **0.56** |
| *Δpde+pPde* | 0.004 ± 0.000 | **2** | 0.005 ± 0.002 | **0.13** |

### The high c-di-AMP strain becomes more prone to UV-induced mutation

After confirming a significant increase in spontaneous mutation rate in the *Δpde* strain, we wanted to estimate the effect of induced mutation with a known mutagenic agent: UV irradiation. Here, we exposed the *M. smegmatis* cells to UV (0.12 mJ/cm^2^), this particular dose was optimized to ensure minimum cell death and maximum mutagenic capacity), grew the cells for 6 days, and plated them on an antibiotic-containing plate to estimate the extent of genome-wide mutations due to UV induced double-strand break and inaccurate repair. In this study, we have included rifampicin (10X MIC) as a selection platform to isolate *rpoB* mutants and as a control, we grew the non-UV exposed cells for 6 days and plated them on the same rifampicin concentration plate to compare the increase in mutation rate specifically after UV exposure. As expected, and as a part of the validation of our assay, we found irrespective of the strain background, *M. smegmatis* WT, *ΔdisA*, and *Δpde* all showed a significant increase in mutation rate upon UV exposure (~40-120 fold) compared to their respective untreated controls under identical assay setup. Comparison in the fold increase of mutation rate showed the highest for the *Δpde*, i.e., 109-fold; which was significantly higher than WT which showed about a 78-fold increase **(Fig. 2A).** The higher fold increase of mutation rate in high c-di-AMP strain reconfirmed our previous observation linking increased c-di-AMP level with enhanced hypermutation phenotype. Microscopic observation of the cells after UV exposure revealed more prominent cell damage and chromosome condensation in the *Δpde* strain **(Fig. 2B).** The lack of cellular filamentation in the *Δpde* strain was also possibly an indication of the absence of RecA-mediated SOS repair. Finally, sequencing of the rifampicin-resistant (rif^R^) clones revealed histidine to tyrosine substitution in the 442^nd^ amino acid of the RpoB protein in representative clones.

**Figure 2.**
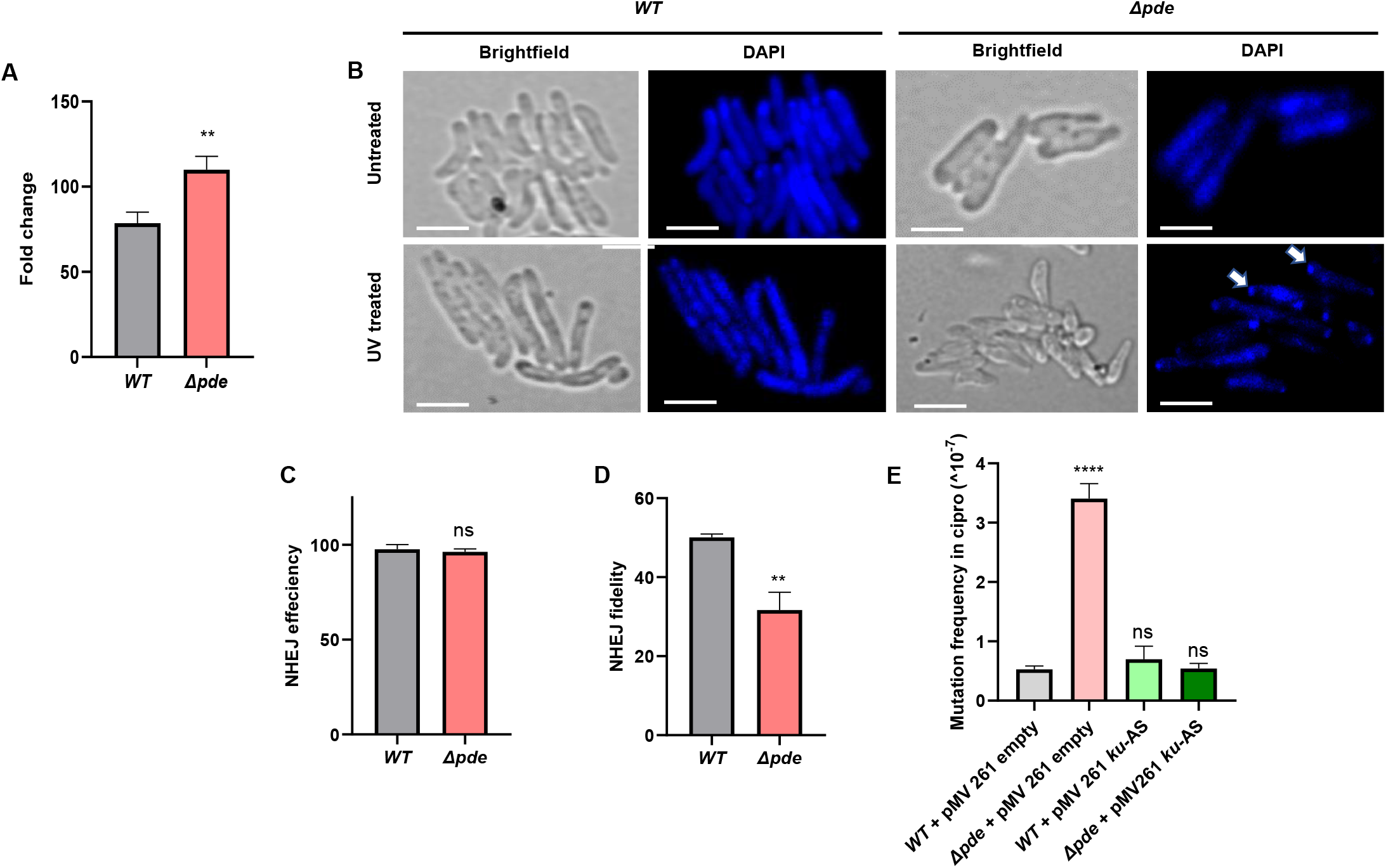
**(A)** Bigger fold change in rifampicin mutation rate was observed for *M. smegmatis Δpde* strain compared to *M. smegmatis* WT strain after UV induced mutagenesis [N=3]. **(B)** Representative microscopic images were shown to compare cell damage, cell filamentation and chromosome disintegration in both *M. smegmatis* WT and *M. smegmatis Δpde* strain after a sublethal dose of UV exposure (0.12 mJ/cm^2^). A lack of cellular filamentation in *Δpde* strain was an indicative absence of RecA-mediated SOS repair in *Δpde* strain. Microscopy scale bars =2 μm.A plasmid-based assay was performed to estimate the NHEJ-driven repair phenomenon. Though **(C)** NHEJ efficiency was comparable between *M. smegmatis* WT and *Δpde* strain [N=3], a significant drop in **(D)** NHEJ repair fidelity (~19%) was observed due to error-prone *lacZ* reannealing [N=3]. **(E)** Knockdown of the principal NHEJ component Ku reversed the hypermutation phenotype in *M. smegmatis Δpde* strain, estimated by resistance frequency calculation against ciprofloxacin [N=3]. All the graphs are plotted using GraphPad Prism8 (N=3), unpaired t-test was used to calculate statistical significance: *** = P < 0.001; ** = P < 0.01; * = P < 0.05; ns= non-significant.

### The *in vivo* NHEJ assay revealed a high degree of error-prone repair in *Δpde* strain

Next, we wanted to understand the underlying molecular mechanism of the enhanced mutation phenotype of the *Δpde* strain. As it has been already highlighted how the RecA enzyme’s function is severely compromised in presence of excess c-di-AMP in the system ^15^, we hypothesized that DNA repair function in such cases becomes largely dependent on the non-homologous end joining (NHEJ) pathway, which is usually considered to be a last-resort, error-prone repair mechanism ^36^. To prove our hypothesis, we adopted a plasmid-based assay to estimate NHEJ-driven repair and compared the accuracy of repair between WT and *Δpde* strain **(Fig.S2a)**. We used the pMV261-*lacZ* reporter plasmid where a functional *lacZ* gene was linearized (single cut) with blunt cutter restriction enzyme SspI, then electroporated the DNA into cells with varying c-di-AMP concentrations and selected on hygromycin+X-gal plate. This ensured simultaneous screening for both antibiotic resistance acquisition (which was possible due to reannealing at the SspI cut site) and *lacZ* enzyme function (which was only possible if the repair is error-free). The total number of blue colonies was counted to determine successful repair events of *lacZ* and the ratio of blue and white colonies estimated the overall repair fidelity of the reporter plasmid. As expected, the uncut reporter plasmid served as a parallel control where all the transformed cells appeared as blue colonies resulting in 100% fidelity. After we made sure the transformation efficiency and NHEJ efficiency is comparable between WT and *Δpde* strains **(Fig. 2C)**, we calculated the percentage of blue colonies on the hygromycin+Xgal plate to estimate faithful repair events. Our data suggested in the *Δpde* strain there was a significant drop in the *lacZ* repair fidelity (~19%) due to error-prone NHEJ repair events **(Fig. 2D)**. Further analysis of repair sites by sequencing revealed the appearance of different base insertions and deletions **(Table S1)** at the SspI restriction site resulting in a frameshift of the amino acid reading frame, which typically occurred due to NHEJ repair, thus reconfirming our hypothesis about the high probability NHEJ driven error-prone repair in high c-di-AMP strain.

### Knockdown of NHEJ component Ku reversed the hypermutation phenotype of *Δpde* strain

As our plasmid-based repair assay revealed a definite role of the NHEJ mechanism in generating resistant mutants, we wanted to check if the inactivation of Ku protein (the principal component of the NHEJ repair pathway) ^37^ could make any difference in *Δpde* strain’s mutational outcome. For this, we adopted the antisense strategy to conditionally knock down the *ku* gene (*MSMEG_55801*) in *Δpde* strain by overexpressing *ku* gene in the reverse orientation from pMV261 vector. Upon checking MIC **(Table S2)** and repeating the mutation frequency determination assay at 10X MIC of ciprofloxacin, we found, as expected, the *Δpde* strain showed a significant increase in resistance frequency compared to WT; but to our surprise, *M. smegmatis Δpde*+ pMV261-*ku*(As) strain showed no increase in the number of resistant mutants compared to WT counterpart of *M. smegmatis*+ pMV261-*ku* As **(Fig. 2E)**. Hence, it was proved that the underlying mechanism of the hyper mutation phenotype of the *Δpde* strain is directly linked to the error-prone NHEJ repair pathway. To check whether the hypermutation phenotype in *Δpde* strain could also be related to the elevated levels of free radicals ^38^, we performed the resistance mutagenesis frequency experiment with thiourea (a known quencher of free radicals) treated cells (100 mM f.c., pretreatment of cells before spreading in ciprofloxacin plate). We could not observe any differences in terms of resistance frequency in thiourea treated *Δpde* cells compared to untreated *Δpde* cells and in both the cases, it showed a significant increase in Resistance frequency (~30 to 50 fold) compared to the respective WT series of thiourea treated and untreated cells **(Fig.S2b)**. This observation nullified the other possibility of the high level of free radicals responsible for generating a high number of resistant mutants in the case of the *Δpde* strain.

### Increased c-di-AMP concentration results in the high probability of non-QRDR mutations in *Δpde* strain

We were further interested to investigate if the predominance of NHEJ repair in *Δpde* strain resulted in a different mutational landscape or not. To check that, we screened approximately 200 cip^R^ clones that arose in the mutation frequency determination experiment for both WT and *Δpde* strains by patching them on low (10X MIC, original concentration at which they were isolated) and high concentrations of ciprofloxacin (60X MIC) plates **(Fig.S3a)**; further on, we described them as category I and category II mutants respectively. To our surprise, we found the majority (~94%) of cip^R^ mutants in *Δpde* strain background were category I mutant (only resistant at to 10X MIC, but does not survive at higher 60X MIC concentration) and only a minority of them (~6%) were able to survive at 60X MIC of ciprofloxacin. A similar comparison in WT cip^R^ did not reveal any striking predominance of category I mutants over category II mutants (~58% vs 42%) **(Fig.3A)**. To validate this unexpected observation, we confirmed the different degrees of ciprofloxacin resistance phenotype of mutants by checking ciprofloxacin minimum inhibitory concentration (MIC) and found the MIC values were increased only by 16-fold for WT category I mutant (MIC 8 μg/ml.) and 128-fold for WT category II mutant (MIC 64 μg/ml.) compared to the WT parental strain (MIC 0.5 μg/ml.) **(Table S3)**. Category II mutants were further tested for cross-resistance against another fluoroquinolone antibiotic ofloxacin and found to have a 64-fold increase in MIC further confirming target gene mutation (data not shown). Next, we sequenced the QRDR region of the representative category II mutants and all of them were identified as having unique point mutations in the 94^th^ amino acid (aspartic acid) of GyrA protein, but no such mutations were detected in any of the category I mutants. At this point, we hypothesized that the category I mutants which showed a lesser degree of MIC modulation and no mutation in the QRDR hotspot region of the *gyrA* gene, could harbor mutations leading to high efflux activity. To prove that, we repeated the ciprofloxacin MIC assay with efflux pump inhibitor CCCP (Carbonyl cyanide m-chlorophenyl hydrazone) at a non-toxic concentration (4 μg/ml.) and indeed we found a decrease in MIC for WT category I mutant (from 8 μg/ml. to 0.125 μg/ml.), but the shift in MIC in presence of CCCP was not observed in WT parental strain (0.5 μg/ml.) **(Table S3)**. This data further indicated the presence of a certain mutation in the efflux pump in category I mutants. Next, we did an efflux assay using EtBr and observed a high degree of efflux activity in WT category I mutant only, but not in WT category II or parental strains **(Fig. 3B)**. Similar observations were recorded for *ΔdisA and Δpde* strains (category I and category II mutants) **(Fig. S3b).** Finally, we sequenced the *lfrR* gene which serves as a repressor of well-known LfrA efflux protein ^39^, and identified unique insertions in all category I mutant strains **(Table S4)**, which have caused an ORF frameshift and complete loss of its repressor activity and thus resulted in the constitutive expression of LfrA. As expected, for category II mutants, we were unable to map any frameshift mutation in the *lfrR* gene, further signifying the effect of target mutation related to the high degree of resistance against ciprofloxacin **(Table S4)**. Though we confirmed unique SNPs in the QRDR region of category II cip^R^ mutants with WT and *ΔdisA* background (4 independent clones for each), for *Δpde* category II mutants it was only detected once (out of 4 clones sent for sequencing). From this data, it could be hypothesized that the c-di-AMP directed error-prone NHEJ repair (which usually results in frameshift mutation) in *Δpde* strain categorically avoided putting mutations in an essential gene *gyrA* or such frameshift mutants with complete loss of GyrA function eventually not got selected in the end.

**Figure 3.**
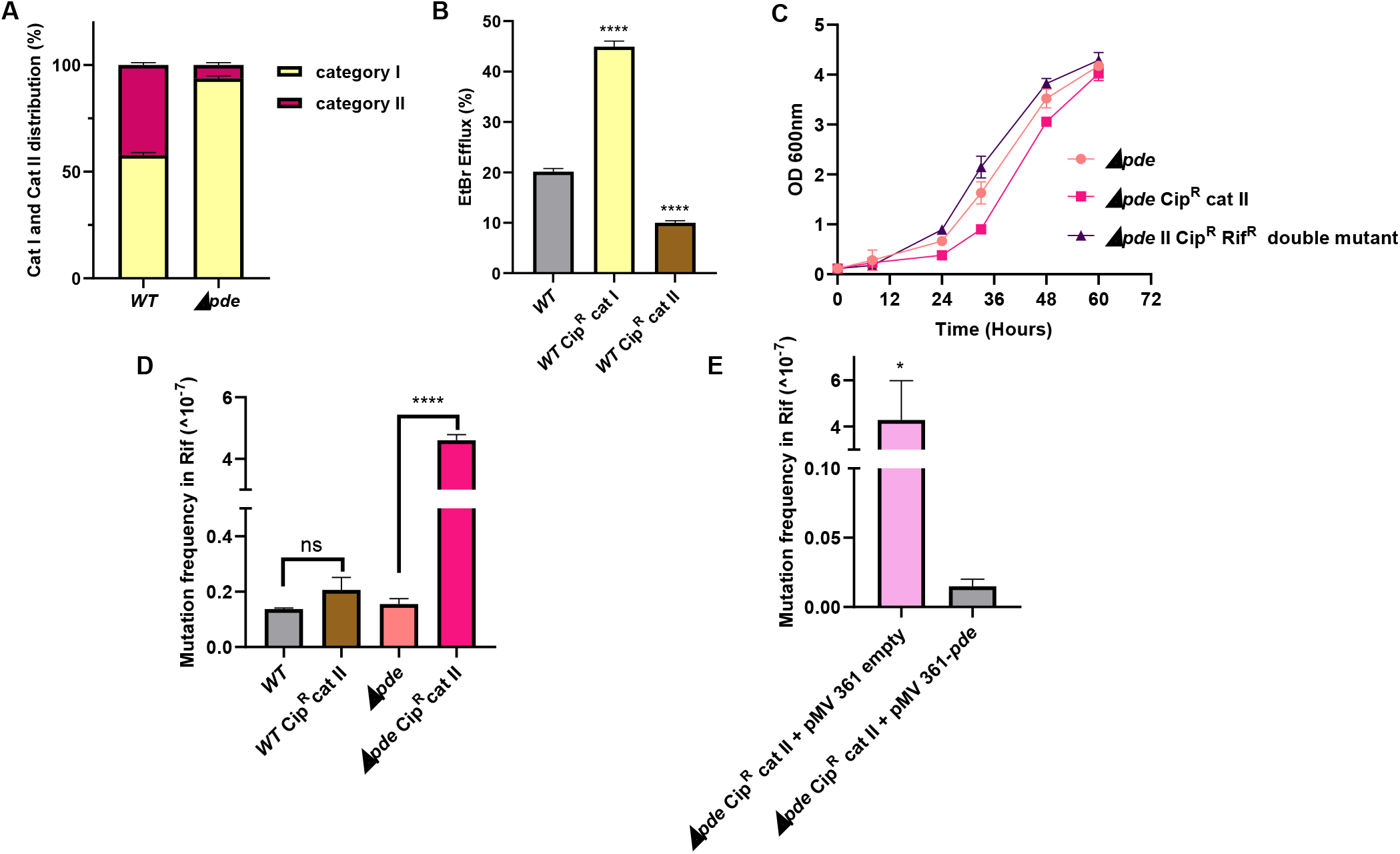
**(A)** Increased c-di-AMP concentration determines the mutational landscape and results in the high probability of non-QRDR (Category-I) mutations in *Δpde* strain, which are only resistant at to 10X MIC, but do not survive at higher 60X MIC of ciprofloxacin. A two-way ANOVA test was performed to check statistical significance [N=2]. **(B)** EtBr efflux assay demonstrated a high degree of efflux activity in category I cip^R^ (non-QRDR) mutant, but not in category II cip^R^ (QRDR SNP) mutant or parental strains [N=3]. **(C)** High fitness cost associated with D94N mutation in the GyrA protein subunit was observed in the case of *Δpde* (cip^R^ category II mutant) when cells had growth deficiency in minimal media and the fitness cost was neutralized to the *M. smegmatis Δpde* parental strain level by acquiring a second compensatory mutation (rif^R^) after prolonged growth in M9 minimal media for 6 days [N=2]. Estimation of the specific role of high intracellular c-di-AMP concentration driving positive epistatic interactions: **(D)** increased rifampicin mutation frequency of a cip^R^ category II mutant to become double mutant (cip^R^, rif^R^) was only favored when intracellular c-di-AMP concentration was high (in *Δpde* background), but not in WT background [N=3] and **(E)** rifampicin mutation frequency dropped in *Δpde* cip^R^ category II mutant when normal physiological level of c-di-AMP was restored with pMV361-*pde* complementation [N=3]. All the graphs are plotted using GraphPad Prism8, unpaired t-test was used to calculate statistical significance: *** = P < 0.001; ** = P < 0.01; * = P < 0.05; ns= non-significant.

### Specific QRDR mutations come with a high fitness cost

Next, we estimated the fitness cost of each QRDR mutation across different strain backgrounds by growing them in stringent conditions such as M9 minimal media, without any ciprofloxacin and checked if the growth profile became different due to the acquisition of mutations in either *gyrA* or *lfrR* genes. In all the cases, *gyrA* 94^th^ amino acid mutation seemed to have a high fitness cost as all category II mutants irrespective of strain backgrounds, grew slowly compared to the corresponding parental strain, and this difference in growth seemed most prominent in the case of *Δpde* strain category II mutant when compared to the parental strain **(Fig. 3C)**, whereas category I mutants of *Δpde* did not reveal any growth deficiency whatsoever. As we had several category II *Δpde* mutants without any QRDR mutation detected, we checked their growth profile in M9 minimal media as well and found that non-QRDR category II mutants had less fitness cost compared to the QRDR category II mutants, though they have the same ciprofloxacin MIC.

### c-di-AMP promotes positive epistatic interaction between resistance genes and results in multi-drug resistance

Since it was clear QRDR mutation usually comes with a high fitness cost, we were further interested to know if a second compensatory mutation could nullify the growth deficiency of a category II mutant. Hence, we grew the parental strain and category II QRDR mutants of both *M. smegmatis* WT and *Δpde* strain for 6 days in M9 media (where they exhibited moderate to high growth deficiency) and screened for the emergence of resistant mutants by plating on different antibiotic rifampicin (10X MIC) plate. To our surprise, we found that only *Δpde* category II mutants formed many colonies on the rifampicin plate and showed a 30-fold increase in mutation frequency compared to *M. smegmatis Δpde* parental strain **(Fig. 3D)**. In the case of WT strain, we could not see such an unanticipated emergence of rifampicin mutants without any pre-exposure to rifampicin. Next, we tested whether due to the acquisition of the second mutation (rifampicin resistance) the fitness cost of the category II mutant was neutralized or not, we checked the growth profile of the double mutant (cip^R^, rif^R^) in M9 minimal media. Indeed, we found the strain has nullified the growth deficiency **(Fig. 3C)** with a second mutation ectopically linked with rifampicin resistance while retaining the first mutation (cip^R^) intact. Subsequently, we checked the rifampicin MIC of the double mutant, we found it was increased by only 16-fold to 32 μg/ml., which is much lesser than the one-step isolated rif^R^ mutant of *Δpde* strain (512 μg/ml.) **(Table S5)**. As expected, *rpoB* gene sequencing of the double mutant did not reveal any point mutations in the RRDR region and it was understood that fitness revival along with low-level rifampicin resistance is not due to RRDR hotspot mutation. Finally, to estimate the specific role of high intracellular c-di-AMP concentration driving positive epistatic interactions, we complemented *Δpde* category II cip^R^ mutant strain with a single copy integrating vector pMV361-*pde*, which was supposed to bring down the high c-di-AMP level to low WT level. First, we checked the growth profile of the *M. smegmatis Δpde* (cip^R^) + pMV361-*pde* strain and found that the growth deficiency of the strain was rectified **(Fig.S3c)**. Upon repeating the same double mutant selection assay in M9 minimal media, we found that in the case of *M. smegmatis Δpde* (cip^R^) + pMV361-*pde*, rifampicin-resistant mutants’ counts were significantly low, similar to *M. smegmatis Δpde* parental strain, whereas the *M. smegmatis Δpde* (cip^R^) + pMV361-Empty strain still generated a high number of double (cip^R^, rif^R^) mutants **(Fig.3E)**. Hence, our data proved that a high intracellular c-di-AMP concentration coupled with a high fitness cost of drug mutation, potentially could drive the evolutionary pattern of the slow-growing strain; specifically, to revive the high fitness cost imposed by the first mutation, cells often spontaneously acquire secondary drug mutation and thus promoting multi-drug tolerance.

### c-di-AMP plays a role in persister cells regrowth by modulating resuscitation-promoting factor gene *rpfA* expression

As our research already pointed out about the definite role of c-di-AMP in resistant mutant evolution, we also wanted to check if increased or lack of c-di-AMP concentration could have a role to play in the persister formation and outgrowth in *M. smegmatis.* In both laboratory and clinical scenarios, antibiotic treatment often gives a biphasic response with a rapid killing phase followed by a plateau which is typically represented by a 0.01-0.001% non-growing persister population ^40^. These antibiotic-selected persister cells often pose a major threat to an effective antibiotic therapy because the persisters could resuscitate into normal growing cells at some point in time after the antibiotic treatment is terminated, resulting in recurrent infection ^41^. First, we began our study by checking persister formation with ciprofloxacin (10XMIC) treatment and we could not find any significant difference (data not shown) between WT, *ΔdisA* and *Δpde* in terms of the percentage of minor survived population after killing ~99.9% of the major population in the culture. Next, we checked the regrowth of persisters with the same strains in presence of a lower concentration of ciprofloxacin (3X MIC), which was extremely relevant in a clinical scenario when the optimum antibiotic concentration could not be maintained due to a variety of reasons. Our data suggested that the regrowth of cells in the complete absence (10X ciprofloxacin treatment followed by washing off the drug) and presence of a lower concentration of ciprofloxacin (3X ciprofloxacin treatment and no washing after 24 hours) significantly decelerated by high c-di-AMP concentration in *Δpde* strain **(Fig.4A, Fig. S4a)**. The complementation strain *M. smegmatis Δpde+* pMV361-*pde* had a similar resuscitation profile to WT **(Fig.4A)**. To further evaluate the direct role of c-di-AMP in the resuscitation of persisters, we used a mutant strain of *M. smegmatis Δpde*, where the *disA* gene function was purposefully abolished by introducing a frameshift mutation in the ORF. As expected, this double mutant strain (*Δpde*, *disA*-out of frame) behaved like WT and further corroborated the previous observation that the high c-di-AMP concentration in *Δpde* strain is specifically responsible for slower resuscitation and it was not due to any polar effect **(Fig. S4b)**. Next, we checked if the slower resuscitation of *Δpde* persister cells could result in more genetic mutants, we estimated the number of ciprofloxacin-resistant genetic mutants (cip^R^) by plating parallelly in ciprofloxacin containing plate. To our surprise, we found a visible increase in cip^R^ mutants in the case of *Δpde* mutant strain which attained a remarkable ~93% mutant population takeover within 24 hours of the regrowth phase, whereas in the case of WT strain it was ~52% of cells became ciprofloxacin mutants **(Fig.4B)**. To know more about the mechanistic insights, we dig into our RNA-seq data and found that the *rpfA* (*MSMEG_5700*) gene was significantly downregulated (transcriptionally) in the *Δpde* strain (log2 fold change -3.0062) and previous studies suggested that the *rpfA* gene encodes for a secreted protein and participated in the resuscitation of non-growing cells ^42^. To our surprise, a detailed search in the prokaryotic riboswitch database (http://ribod.iiserkol.ac.in/genome_search2.php?id=NC_008596.1) revealed the presence of an *ydaO-yuaA* class of riboswitch just upstream of the *rpfA* gene. The *ydaO-yuaA* has been previously characterized as a c-di-AMP responsive riboswitch in *Bacillus subtilis* and other bacteria ^43^. To check if there is any possible interaction between c-di-AMP and riboswitch element upstream of *rpfA*, we designed a transcriptional fusion by fusing 5’-UTR region of *rpfA* (434 bp) to promoter-less GFP in pMN406 plasmid. Next, we used this reporter plasmid in different strain backgrounds to check GFP fluorescence (as a measure of promoter induction) during the resuscitation phase of persisters. Indeed, we found a significantly slower induction of the *PrpfA* in *Δpde* background when the minor population of antibiotic tolerant cells was allowed to regrow in fresh media after 15 hours of ciprofloxacin treatment and washing off the drug. As a control, we used the double mutant strain (*Δpde, disA*-out of frame), which behaved like WT in terms of *PrpfA* induction under identical conditions **(Fig.4C)**. This important observation described the molecular basis of the high c-di-AMP driven slower resuscitation of persisters in *M. smegmatis.*

**Figure 4.**
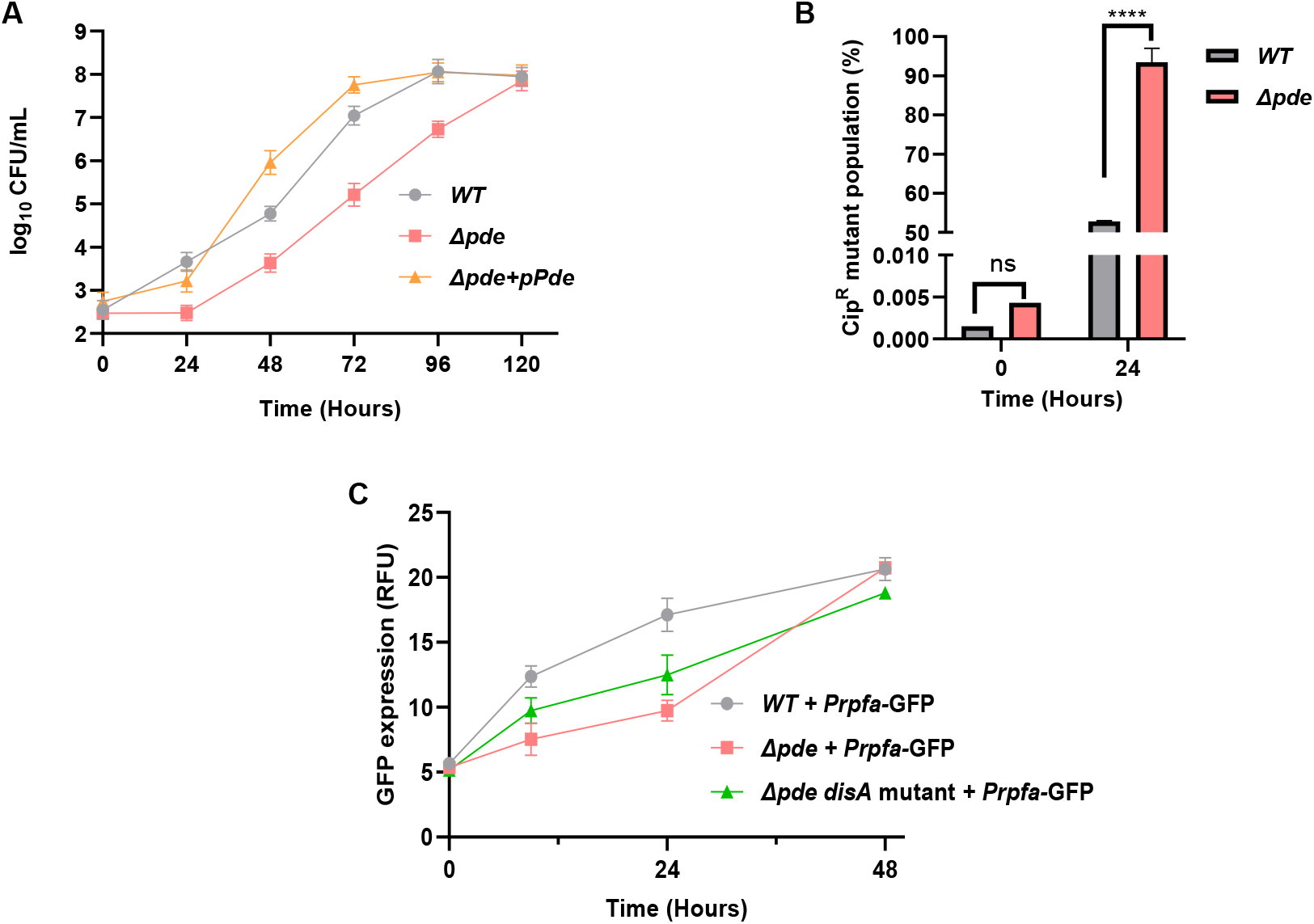
**(A)** Regrowth of persisters (after 10X ciprofloxacin treatment and subsequent washing of the drug) was significantly decelerated by high c-di-AMP concentration in *Δpde* strain, unlike WT and the complementation strain (*Δpde+* pMV361-*pde*) [N=3]. **(B)** Slower resuscitation of *Δpde* persister cells results in a rapid enrichment of genetic mutants, around 93% of mutant population enrichment happened in *Δpde* strain within 24 hours of the regrowth phase, whereas in the case of WT strain ~52% of cells became ciprofloxacin mutants [N=3]. **(C)** Significantly slower induction of the *P_rpfA_* (as a GFP readout) in *Δpde* background confirms the molecular basis of slower persisters regrowth [N=3], when the c-di-AMP level was normalized in the same strain by a *disA* gene frameshift mutation, *PrpfA* induction was almost restored to the WT level. All the graphs are plotted using GraphPad Prism8, unpaired t-test was used to calculate statistical significance: *** = P < 0.001; ** = P < 0.01; * = P < 0.05; ns= non-significant.

## Discussion

Bacterial second messengers have been shown to play important roles in the cellular physiology of different bacteria. In a previous study, we thoroughly studied the role of c-di-AMP in *M. smegmatis* starting from basic phenotypes modulation to stress response and antibiotic sensitivity. As a continuation of that study, here we were particularly interested to explore the role of c-di-AMP in generating antibiotic tolerant cells. We used two different deletion mutants of *M. smegmatis* with varying c-di-AMP concentrations and confirmed that both spontaneous mutation frequency and mutation rate got increased by several folds when the intracellular c-di-AMP level is higher than usual. To confirm the direct involvement of high c-di-AMP, we included *Δpde* complementation (*M. smegmatis Δpde+* pMV361-Pde) and found that the hypermutation phenotype is reversed to the normal WT level.

Next, we found that if cells were treated with a DNA-damaging agent (mutagen), the possibility of genome-wide mutation is directly proportional to the c-di-AMP concentration, which was hypothesized to be linked with the lack or absence of the RecA mediated Homology directed repair (HDR) mechanism. In such cases, *M. smegmatis* cells tend to become largely dependent on the non-homologous end joining (NHEJ) pathway, which is fallible and can give rise to unwanted mutations. Our plasmid-based *in vivo* NHEJ assay result indeed pointed towards high mutation probability in *Δpde* strain as evidenced by a significant decrease in repair fidelity of the reporter gene *lacZ.* Conditional knockdown of the main component of the NHEJ pathway, Ku protein reverted the hypermutation phenotype in *Δpde* strain, which was helpful to precisely understanding the molecular basis of the phenotype. After understanding the molecular mechanism behind the high mutation phenotype, we did a mutant screening exercise of cip^R^ mutants and discovered a predominantly high proportion of clones without QRDR mutations in *Δpde* strain, and often acquiring mutations in efflux pump repressor *lfrR*, which should be sufficient to grow at 10X MIC concentration of the drug. We hypothesize that the rarity of QRDR mutations in high c-di-AMP strain is related to the high degree of error-prone NHEJ repair-driven loss of GyrA function. Occasionally, *Δpde* strain harbored a QRDR mutation (D94N) but that caused a considerable fitness loss of the strain and that could be also a reason for the non-selection of QRDR mutants.

Next, we checked if the hypermutant *Δpde* strain could rescue the fitness defect posed by a single point mutation in the QRDR region of the *gyrA* gene. We found that while growing under stringent conditions for a long time where the fitness cost seemed to be very high, the high c-di-AMP strain tends to put a secondary mutation somewhere else in the genome to revive the fitness back. A further screening revealed that the cip^R^ mutant could evolve to become resistant to rifampicin (with significantly high resistant frequency compared to the *Δpde* parental strain) and thus neutralized the high fitness cost posed by the first mutation. In, WT and *ΔdisA* QRDR mutant strains (which also had moderate growth deficiency), we could not observe such single mutant (cip^R^) to double mutant (cip^R^, rif^R^) transition and hence pointed out the relevant role of c-di-AMP in mediating positive epistatic interaction between two resistance genes. The fact that complementation of *Δpde* strain with single copy of *pde* gene (pMV361-*pde*) reversed the epistasis phenomenon pointed out the critical role of c-di-AMP in the process.

In the final section of the study, we assessed how the c-di-AMP concentration plays a role in persister cells regrowth kinetics. The *Δpde* strain always grew slow during the regrowth phase following ciprofloxacin treatment and by multiple means when we neutralized the high c-di-AMP concentration in the *Δpde* strain, we observed uninhibited resuscitation similar to WT. This made us curious to discover if c-di-AMP was regulating some protein responsible for resuscitation, our RNA-seq data and subsequent bioinformatic search indeed identified a particular gene *rpfA*, which seemed to be transcriptionally downregulated by c-d-AMP because of a putative riboswitch element in the 5’-UTR region. By using a GFP-based transcriptional fusion construct we indeed proved that *PrpfA* had significantly reduced activity under high c-di-AMP concentration.

All in all, our model suggested that c-di-AMP plays a key role in the resistant mutant generation and promotes multidrug tolerance in *M. smegmatis* **(Fig. 5)** as well as persisters regrowth kinetics. In both cases, we have identified the underlying molecular mechanisms to elucidate the role of c-di-AMP and associated downstream components to drive antibiotic resistance. So far, as per our knowledge, there was no published report describing the direct involvement of a signaling messenger molecule to promote the generation of resistant mutants, especially promoting multi-drug tolerance via positive epistatic interaction between two different antibiotics, which we think is the major highlighting point of the study and extremely relevant for understanding the multi-drug tolerance phenotypes of mycobacteria in clinical perspective.

**Figure 5.**
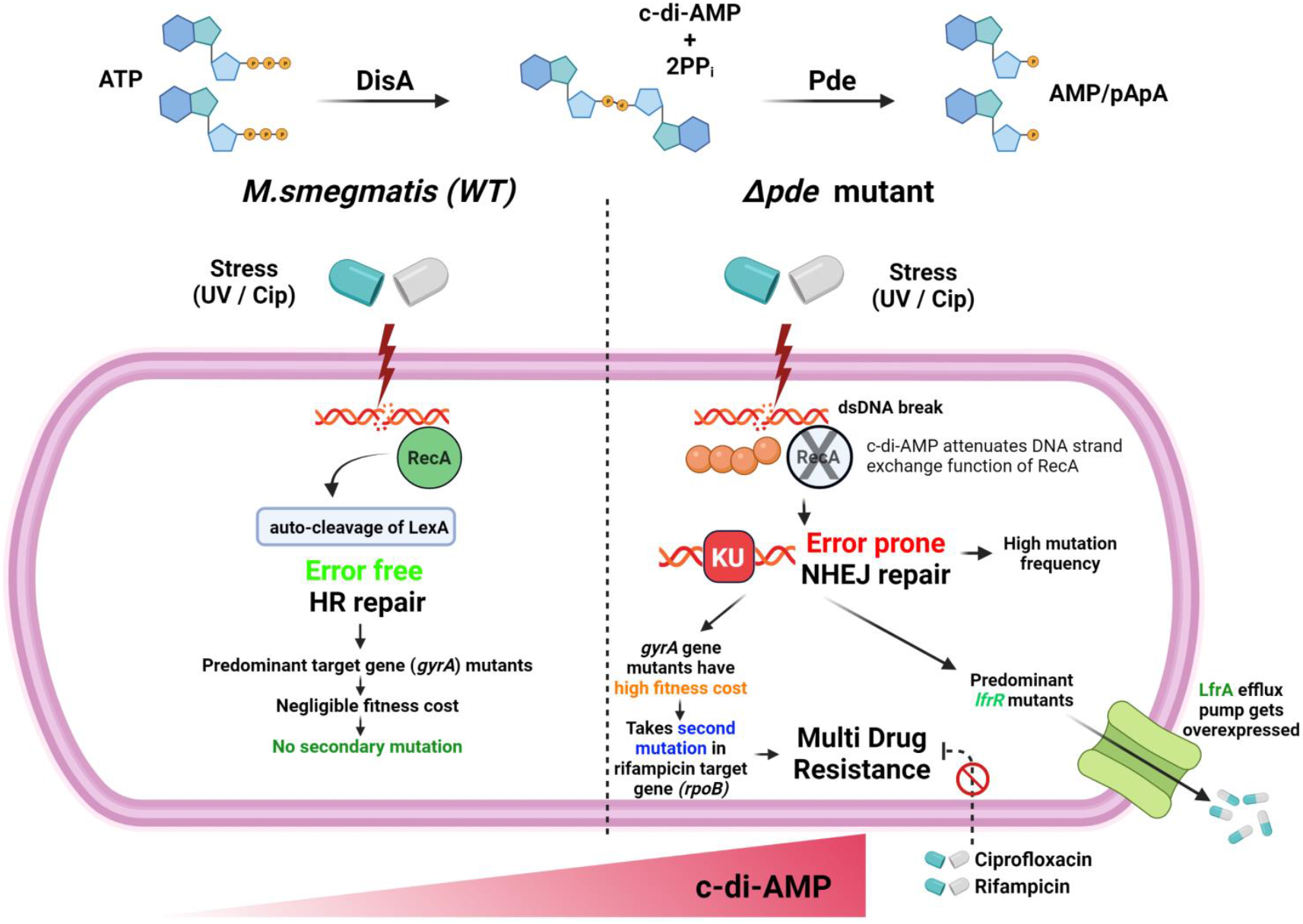
Schematic representation of our proposed model showing how high c-di-AMP concentration in *M. smegmatis* drives error-prone DNA repair which results in a significant change in the mutational landscape and fitness of a cell and finally promotes multi-drug resistance (figure created with the paid license from www.biorender.com).

## Acknowledgment

AG thanks the Ramalingaswami Re-entry Fellowship and the Department of Biotechnology (DBT), Government of India, for funding this work (BT/RLF/Re-entry/31/2017). AP acknowledges the Department of Biotechnology (DBT), Government of India for his fellowship. We thank Prof. Dipankar Chatterji, MBU, IIsc for valuable feedback on the work. AP and AG contributed to the conception and design of the study. AP and AG performed the experiments. AP and AG participated in data analysis and interpretation. AP and AG wrote the manuscript. The authors declare no conflict of interest.

