## Supplemental figures and tables for "Understanding the role of c-di-AMP signaling in determining antibiotic tolerance in *Mycobacterium smegmatis*: generation of resistant mutants and regrowth of persisters"

### Supplementary material

#### Figure S1

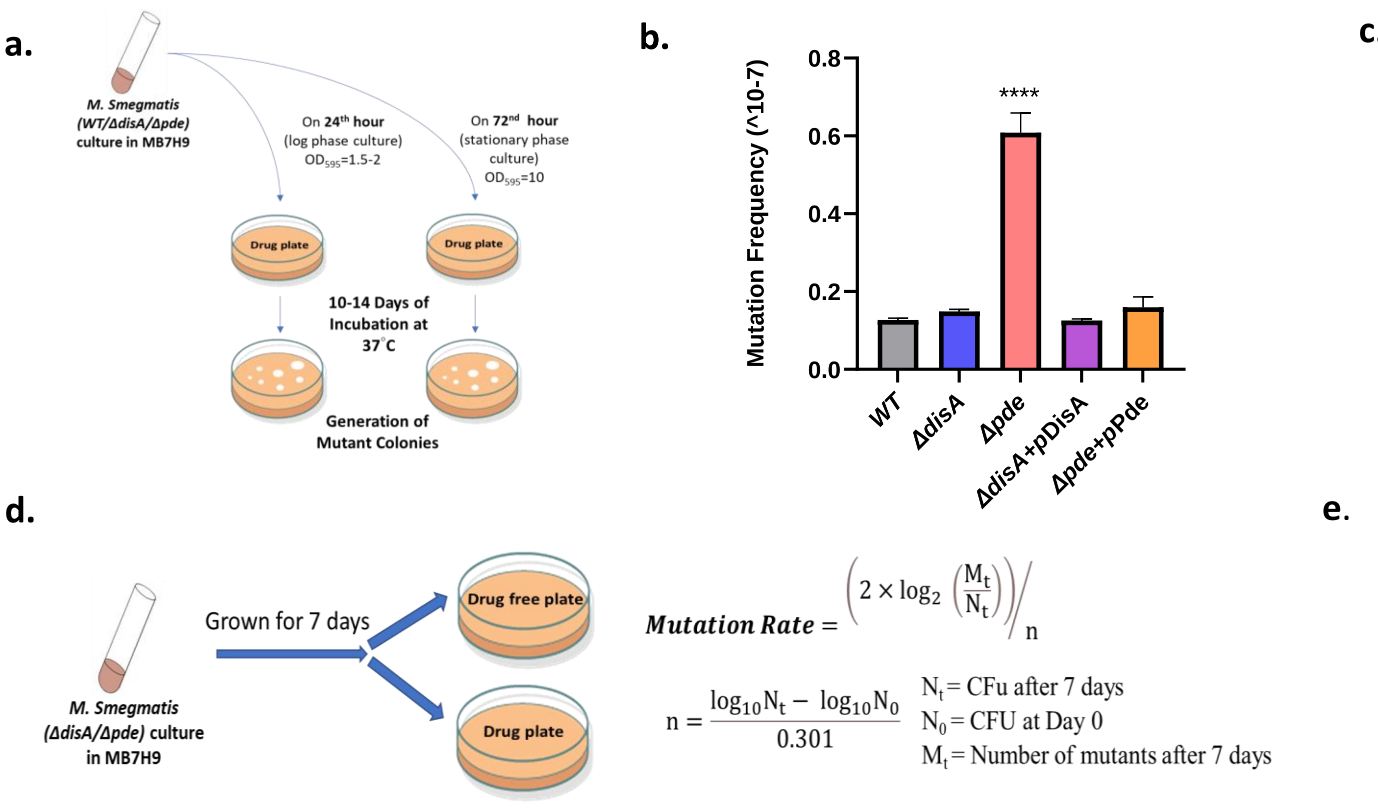


$$\textit{Mutation Rate} = \left(2 \times \log_2\left(\frac{M_t}{N_t}\right)\right)\Big/ n$$

$$n = \frac{\log_{10} N_t - \log_{10} N_0}{0.301}$$

$N_t$ = CFu after 7 days
$N_0$ = CFU at Day 0
$M_t$ = Number of mutants after 7 days

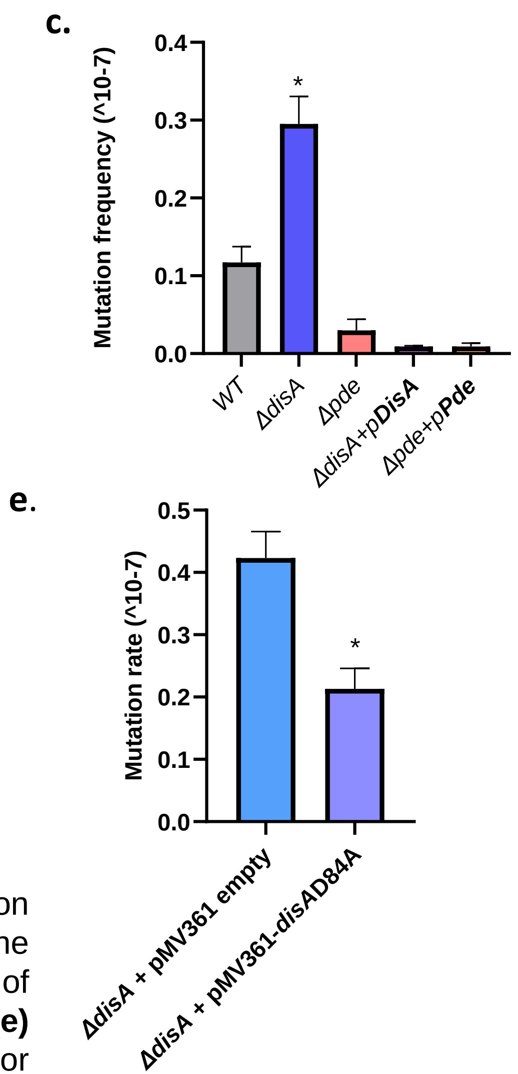


**Figure S1:** **(a)** Schematic illustration of the experimental setup to enumerate mutation frequency; Mutation frequency determination using exponentially grown cultures of the respective strains for **(b)** ciprofloxacin and **(c)** rifampicin [N=3]. **(d)** Schematic illustration of the experimental setup to enumerate mutation rate along with the calculation formula; **(e)** Mutation rate calculation for *ΔdisA* strain when complemented with either pMV361-Empty or pMV361-*disA(D84A)* vector.

### Figure S2

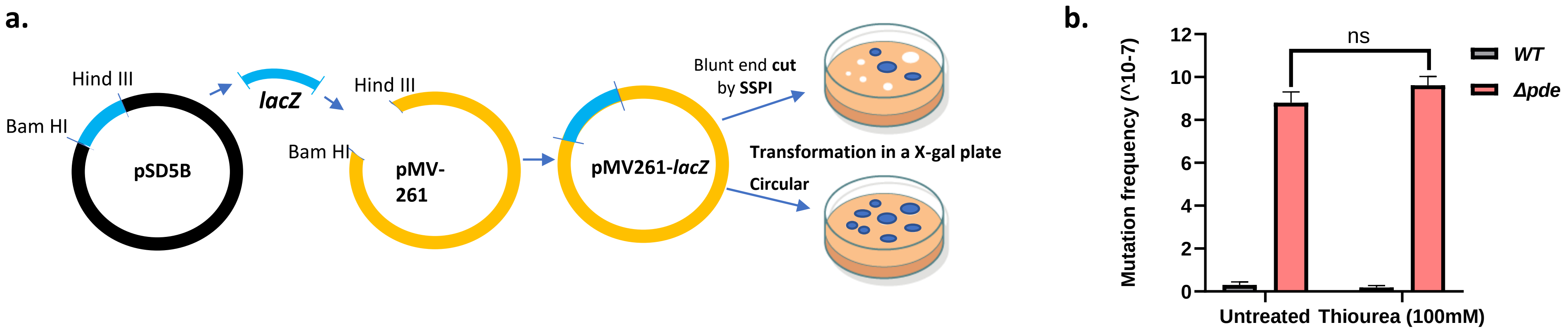


**Figure S2: (a)** Schematic representation depicting the construction of the *lacZ*-reporter plasmid and NHEJ-driven plasmid repair and fidelity assay; **(b)** Ciprofloxacin mutation frequency determination of the respective strains in the presence and absence of thiourea (100mM).

#### Figure S3

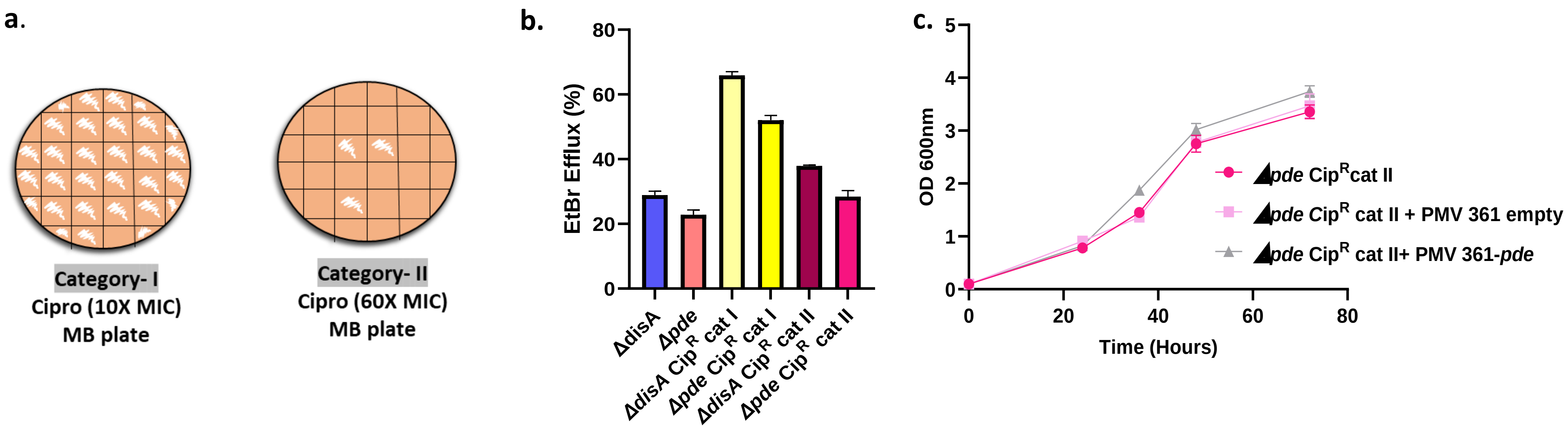


**Figure S3: (a)** Schematic illustration of the screening for Category-I and Category-II cipR mutants; **(b)** EtBr efflux assay for different categories of *M. smegmatis ΔdisA* and *M. smegmatis Δpde* cipR mutants; **(c)** Minimal media growth curve depicting fitness cost of respective mutant strains.

#### Figure S4

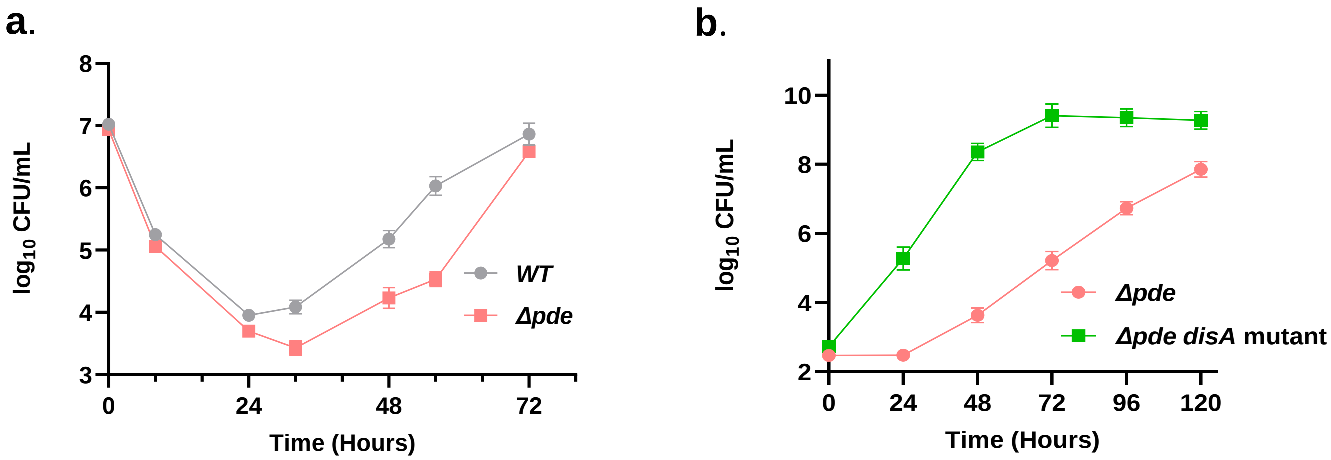


**Figure S4:** Regrowth of persisters **(a)** in presence of a low concentration of ciprofloxacin (3X MIC) for WT and *Δpde* strains and **(b)** in the complete absence of ciprofloxacin (10X treatment followed by washing off the drug) for *Δpde, disA-fs* strains.

#### Table S1

| Sl No. | *Δpde LacZ* mutant (SSPI cut site - AATATT) | |
|---|---|---|
| **1** | **AATTATT** | T insertion |
| **2** | **AATTATT** | T insertion |
| **3** | **A--ATT** | AT deletion |
| **4** | **AATTATT** | T insertion |
| **5** | **AATAATT** | A insertion |

**Table S1:** Sequencing confirmation of frameshift mutations at the repair sites of *Δpde* + pMV261-*lacZ* white clones

#### Table S2

| **Strain** | **MIC (µg/ml)** |
|---|---|
| | **Ciprofloxacin** |
| ***WT*** **+ pMV261 empty** | 0.25 |
| ***Δpde*** **+ pMV261 empty** | 0.125 |
| ***WT*** **+ pMV261** ***ku*****-AS** | 0.25 |
| ***Δpde*** **+ pMV261** ***ku*****-AS** | 0.0625 |

**Table S2:** Ciprofloxacin MIC values of the respective strains

#### Table S3

| Strain | MIC (µg/ml) Ciprofloxacin |
|---|---|
| **WT** | 0.25 |
| **WT cip$^R$ cat I** | 8 |
| **WT cip$^R$ cat II** | 64 |
| ***ΔdisA*** | 0.25 |
| ***ΔdisA* cip$^R$ cat I** | 8 |
| ***ΔdisA* cip$^R$ cat II** | 64 |
| **Δpde** | 0.125 |
| ***Δpde* cip$^R$ cat I** | 4 |
| ***Δpde* cip$^R$ cat II** | 32 |
| **WT cip$^R$ cat I + CCCP (5µg/mL)** | >0.125 |
| **WT cip$^R$ cat II + CCCP** | 16 |

**Table S3:** Ciprofloxacin MIC values of the parental and mutant strains

#### Table S4

| Sl No. | Strain | Mutant sequencing: QRDR of *gyrA* *(MSMEG_0006)* | Mutant sequencing: *lfrR* *(MSMEG_6223)* |
|---|---|---|---|
| 1 | ***WT* cip$^R$ cat I** | No mutation | L59fs |
| 2 | ***WT* cip$^R$ cat II** | D94Y (2)<br>D94G (2) | NA |
| 3 | ***ΔdisA* cip$^R$ cat I** | No mutation | H29fs |
| 4 | ***ΔdisA* cip$^R$ cat II** | D94G (2)<br>D94Y (1) | NA |
| 5 | ***Δpde* cip$^R$ cat I** | No mutation | P167fs |
| 6 | ***Δpde* cip$^R$ cat II** | D94N (1) | NA |

**Table S4:** Sequencing confirmations of different amino acid substitutions and frameshifts in *gyrA* and *lfrR* genes of the cipR mutants

### Table S5

| Strain | MIC (µg/ml) |
|---|---|
| | Rifampicin |
| ***Δpde*** | 2 |
| ***Δpde* Rif$^{R}$ mutant** | 512 |
| ***Δpde* II Cip$^{R}$ Rif$^{R}$ double mutant** | 32 |

**Table S5:** Rifampicin MIC values of the parental and mutant strains

### Table S6

| **Strains** | **Description** | **Source/ Reference** |
|---|---|---|
| *M. smegmatis* Wild type(WT) | *M. smegmatis* $mc^2$ 155 | Laboratory stock |
| *M. smegmatis ΔdisA* | *M. smegmatis* $mc^2$ 155 strain knock out for disA ; $Kan^R$ | Ref.1 |
| *M. smegmatis Δpde* | *M. smegmatis* $mc^2$ 155 *strain knock out for pde ;* $Kan^R$ | Ref.1 |
| *M. smegmatis ΔdisA* +*p*DisA | *M. smegmatis ΔdisA* strain containing disA cloned in pMV361 vector; $Hyg^R$,$Kan^R$ | Ref.1 |
| *M. smegmatis Δpde* +*p*Pde | *M. smegmatis Δpde* strain containing pde cloned in pMV361 vector; $Hyg^R$,$Kan^R$ | Ref.1 |
| *M. smegmatis ΔdisA* +*disA* D84A | *M. smegmatis ΔdisA* strain containing disA (D84A) cloned in pMV361 vector; $Hyg^R$,$Kan^R$ | Ref.1 |
| *M. smegmatis ΔdisA* + pMV361 empty | *M. smegmatis ΔdisA* strain containing pMV361 empty vector; $Hyg^R$,$Kan^R$ | Ref.1 |
| *M. smegmatis WT* +pMV261 empty | *M. smegmatis* WT strain containing pMV261 empty vector; $Hyg^R$,$Kan^R$ | This study |
| *M. smegmatis Δpde* +pMV261 empty | *M. smegmatis* WT strain containing pMV261 empty vector; $Hyg^R$,$Kan^R$ | This study |
| *M. smegmatis WT* + pMV261 *ku*-AS | *ku*-anti sense knockdown construct *(MSMEG_5580)* in pMV261:hyg vector transformed into *M. smegmatis* WT | This study |
| *M. smegmatis Δpde* + pMV261 *ku*-AS | *ku*-anti sense knockdown construct *(MSMEG_5580)* in pMV261:hyg vector transformed into *M. smegmatis Δpde* | This study |

**Table S6:** Bacterial strains used in the study

#### Table S6 (cont..)

| **Strains** | **Description** | **Source/ Reference** |
|---|---|---|
| *M. smegmatis Δpde disA* mutant | *M. smegmatis Δpde* with a frameshift mutation in *disA* gene | This study |
| *M. smegmatis* WT $cip^R$ cat I | Ciprofloxacin resistant mutant of *M. smegmatis* WT having mutation in *lfrR* gene | This study |
| *M. smegmatis* WT $cip^R$ cat II | Ciprofloxacin resistant mutant of *M. smegmatis* WT having mutation in *gyrA* gene | This study |
| *M. smegmatis* Δ*disA* $cip^R$ cat I | Ciprofloxacin resistant mutant of *M. smegmatis* Δ*disA* having mutation in *lfrR* gene | This study |
| *M. smegmatis* Δ*disA* $cip^R$ cat II | Ciprofloxacin resistant mutant of *M. smegmatis* Δ*disA* having mutation in *gyrA* gene | This study |
| *M. smegmatis* Δ*pde* $cip^R$ cat I | Ciprofloxacin resistant mutant of *M. smegmatis* Δ*pde* having mutation in *lfrR* gene | This study |
| *M. smegmatis* Δ*pde* $cip^R$ cat II | Ciprofloxacin resistant mutant of *M. smegmatis* Δ*pde* having mutation in *gyrA* gene | This study |
| *M. smegmatis* Δ*pde* $cip^R$ cat II $rif^R$ double mutant | Mutant of *M. smegmatis* Δ*pde* having resistance to ciprofloxacin and rifampicin | This study |
| *M. Smegmatis Δpde* $rif^R$ mutant | Rifampicin resistant mutant of *M. smegmatis* Δ*pde* having mutation in *rpoB* gene | This study |
| WT $cip^R$ cat II + pMV 361 empty | *M. smegmatis* WT $Cip^R$ cat II having pMV361 vector; $Hyg^R$ | This study |
| *WT* $cip^R$ cat II + pMV 361 *pde* | *M. smegmatis* WT $Cip^R$ cat II having pMV361 vector; $Hyg^R$ | This study |
| *M. smegmatis* WT + *Prpfa*-GFP | *M. smegmatis* WT with promoter fusion construct (434us) rpfa- $pMN406_{\Delta imyc}$ ; $Hyg^R$ | This study |
| *M. smegmatis Δpde* + *Prpfa*-GFP | *M. smegmatis Δpde* with promoter fusion construct (434us) rpfa- $pMN406_{\Delta imyc}$ ; $Hyg^R$ | This study |
| *M. smegmatis Δpde disA* mutant + *Prpfa*-GFP | *M. smegmatis Δpde disA* mutant with promoter fusion construct (434us) *(rpfa- $pMN406_{\Delta imyc}$ ;* $Hyg^R$ | This study |
| Δ*pde* + pMV261 *lacZ* | *M. smegmatis Δpde* with *lacZ* gene cloned in pMV261 vector *;* $Hyg^R$ | This study |

**Table S6:** Bacterial strains used in the study

#### Table S7

| Name | Sequence (5’ to 3’) | Description |
|---|---|---|
| lacZ_PMV261FW_hindIII | AGCT*AAGCTT*TCCGGAGGAATCACTTCCAT | To construct the NHEJ reporter vector |
| lacZ_PMV261REV_hindIII | TCAG*AAGCTT*TTATTTTTGACACCAGACCA | To construct the NHEJ reporter vector |
| Ku_AS_REV_HindIII | ATCGAAGCTTATGAACCGTGCGGTACGCCA | To construct knock down of *ku* gene |
| Ku_AS_FW_BamHI | ACTAGGATCCCTACGACTTCTTCGCAGCTG | To construct knock down of *ku* gene |
| Ms_QRDR FW | GACCGACATCGGTGGGTTGC | To amplify QRDR region of *gyrA* |
| Ms_QRDR Rev | AATCGACTGTCTCCTCGTCG | To amplify QRDR region of *gyrA* |
| lfrR_fw | ACCGCCTCCGGCGGTGCCGGCT | To amplify QRDR region of *lfrR and also u*sed for sequencing of *lfrR* |
| lfrR_rev | GCCACCCAGGCGCGCGTGGGCG | To amplify QRDR region of *lfrR* |
| rrdr_fw | GTCGTCTGCGCACCGTCGG | To amplify RRDR region of *rpoB and also u*sed for sequencing of RRDR |
| rrdr_rev | CTCGATGAAGCCGAACGGG | To amplify RRDR region of *rpoB* |
| gyrA_fw | ATGACTGATACGACGCTGCCGC | Used for sequencing of *gyrA* |
| lacZ_mid_rev_2 | GCCTTCATACTGCACCGGGCG | Used for sequencing of *lacZ* |
| disa_mid_rev | GGTCATCACGTCGCGCAGCG | Used for sequencing of *disA* |
| rpfa_434bpus _FW_Xba1 | AGCAGTCTAGACGTGAGATACGTCACATC | Used for making *PrpfA*-GFP transcriptional fusion construct |
| rpfa_434bpus_Rev_Sph1 | TCATAGCATGCAAGCGTCGAGGTCCTCTC | Used for making *PrpfA*-GFP transcriptional fusion construct |

**Table S7:** Primers used in the study
